# ChemPert: mapping between chemical perturbation and transcriptional response for non-cancer cells

**DOI:** 10.1101/2022.04.29.490084

**Authors:** Menglin Zheng, Satoshi Okawa, Miren Bravo, Fei Chen, María-Luz Martínez-Chantar, Antonio del Sol

**Affiliations:** Luxembourg Centre for Systems Biomedicine (LCSB), University of Luxembourg, 6 Avenue du Swing, Esch-sur-Alzette, L-4367 Belvaux, Luxembourg; Liver Disease Laboratory, Center for Cooperative Research in Biosciences (CIC bioGUNE), Basque Research and Technology Alliance (BRTA), Bizkaia Technology Park, Derio, Spain; Centro de Investigación Biomédica en Red de Enfermedades Hepáticas y Digestivas (CIBERehd), 48160, Bizkaia, Spain; German Research Center for Artificial Intelligence (DFKI), 66123, Saarbrücken, Germany; CIC bioGUNE-BRTA (Basque Research and Technology Alliance), Bizkaia Technology Park, 801 Building, 48160 Derio, Spain; IKERBASQUE, Basque Foundation for Science, Bilbao 48013, Spain

## Abstract

Prior knowledge of perturbation data can significantly assist in inferring the relationship between chemical perturbations and their specific transcriptional response. However, current databases mostly contain cancer cell lines, which are unsuitable for the aforementioned inference in non-cancer cells. Here we present ChemPert (https://chempert.uni.lu/), a database consisting of 82270 transcriptional signatures across 167 non-cancer cell types, enabling more accurate predictions of perturbation responses and drugs compared to cancer databases in non-cancer cells. In particular, ChemPert correctly predicted drug effects for treating non-alcoholic steatohepatitis and novel drugs for osteoarthritis. Overall, ChemPert provides a valuable resource for drug discovery in non-cancer diseases.

## Introduction

The inference of the relationship between chemical perturbations and their specific transcriptional response has wide biological and clinical relevance, such as drug discovery. However, the inference of such relationship using computational models of signal transduction remains a challenge, as they require data for different molecular regulatory layers, such as phosphoproteomics data, which are not widely available. On the other hand, the analysis of transcriptomics changes before and after perturbations enables us to directly map the chemical perturbations to their response genes. However, a major limitation is that such transcriptional changes (i.e., transcriptional signatures) are usually cell specific and need to be generated for each cell type of interest, necessitating a large compendium of gene expression profiles for large-scale drug screening.

In an effort to address this important challenge, the Connectivity Map (CMap) project and more recently, the LINCS L1000 project, have collected gene expression profiles for thousands of perturbagens at different time points and doses in different cell lines (1,2). These resources have been successfully employed for various studies (3,4). However, the majority of the gene expression profiles in these compendia consist of cancer cell lines, which are known to exhibit significantly distinct signal transduction pathways and gene regulatory networks from non-cancer cells (5). For this reason, the gene expression profiles in these resources are not suitable for mapping signaling perturbations to transcriptional responses in non-cancer cells.

In this study we present ChemPert (https://chempert.uni.lu/), a comprehensive compendium of manually curated transcriptional signatures derived solely from non-cancer cell perturbation datasets, combined with a tool that allows users to predict either the transcriptional responses of perturbations or chemical compounds targeting desired sets of transcription factors. The chemical perturbations in ChemPert are denoted as perturbagens, which include both chemical and biological agents such as small molecules, drugs, cytokines and growth factors. ChemPert consists of 82270 transcriptional signatures of 167 unique non-cancer cell types perturbed with 2566 unique perturbagens. Unlike the existing methods that predict chemical compounds directly from a database (1,2), ChemPert first predicts signaling proteins and then identifies potential perturbagens targeting these proteins. This approach allows for the identification of novel perturbagens that are not contained in the collected transcriptional compendium.

We show that predictions generated for non-cancer cells when using ChemPert database were significantly more accurate than those based on cancer databases, underscoring the importance of non-cancer cell perturbation datasets collected in this study. Our benchmarking also reveals that considering initial cell states in addition to perturbagen similarity for TF response prediction results in significantly higher predictive accuracy than using perturbagen similarity alone. In particular, the application of ChemPert to non-alcoholic steatohepatitis (NASH) models generated novel predictions of the differential TF responses of chemical drugs, which were consistent with their reported functional effects on different stages of NASH. Furthermore, the application of ChemPert to osteoarthritis (OA) successfully recapitulated chemical compounds for partial OA therapies and predicted novel ones. Notably, no effective pharmacologic treatments are currently available for OA and the predicted perturbagens constitute potential novel therapeutics that could be further experimentally validated.

Overall, ChemPert provides a comprehensive non-cancer cell perturbation compendium and facilitates future *in silico* predictions of perturbation response and chemical compound discovery for inducing desired effects on non-cancer cells.

## Results

### Overview of ChemPert

In order to infer the relationship between the signalling perturbation and downstream transcriptional responses, we exhaustively collected and compiled transcriptome profiles of chemical perturbations applied solely on non-cancer cells from public resources (Figure 1A). Based on the database, we developed a tool that allows users to predict either the transcriptional responses of perturbations or chemical compounds targeting desired sets of TFs (Figure 1B). The database consisting of 82270 transcriptional signatures derived from 2566 unique perturbagens across 167 unique normal cell types/lines/tissues (Figure 2A). The datasets covered 2132 unique TFs, in both activation (up) and inhibition (down) directions with no significant bias towards either of them (Figure 2B). More than half of the perturbagens (^~^65%) have frequency not larger than 20 (Figure 2C) and majority of the perturbagens (^~^98%) in the ChemPert database have duration not larger than 24 hours (Figure 2D). In addition, we also collected and integrated the protein targets and corresponding effects (activation, inhibition or unknown) of 57818 chemical compounds.

**Figure 1.**
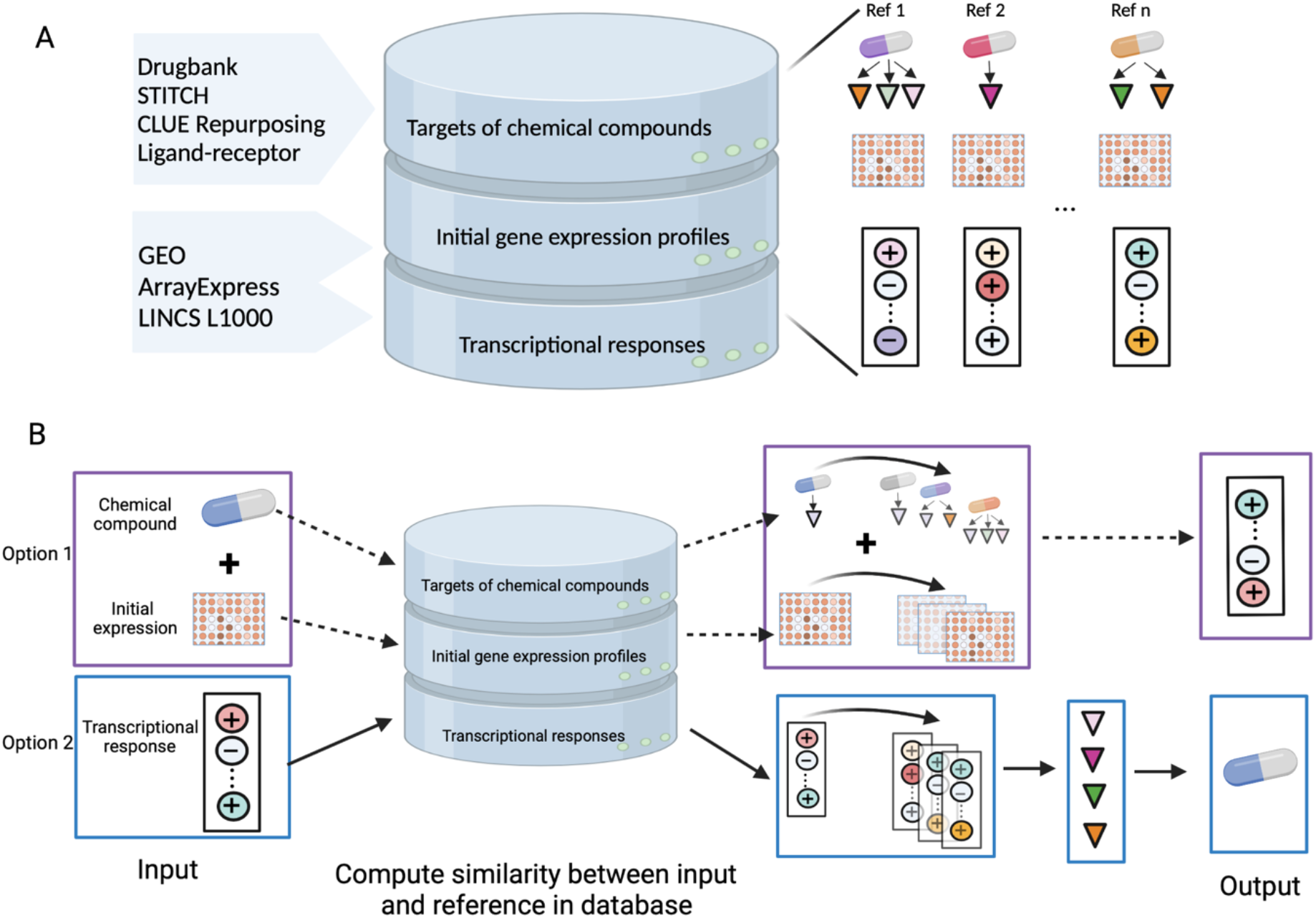
Schematic outline of ChemPert. (**A**) The sources and three main components of ChemPert database. (**B**) Illustration of bulit-in algorithms in ChemPert. One option for predicting the TF responses given the perturbagen and expression profile of initial cellular state (Option 1) and the other for predicting perturbagens that induce desired transcriptional response (Option 2).

**Figure 2.**
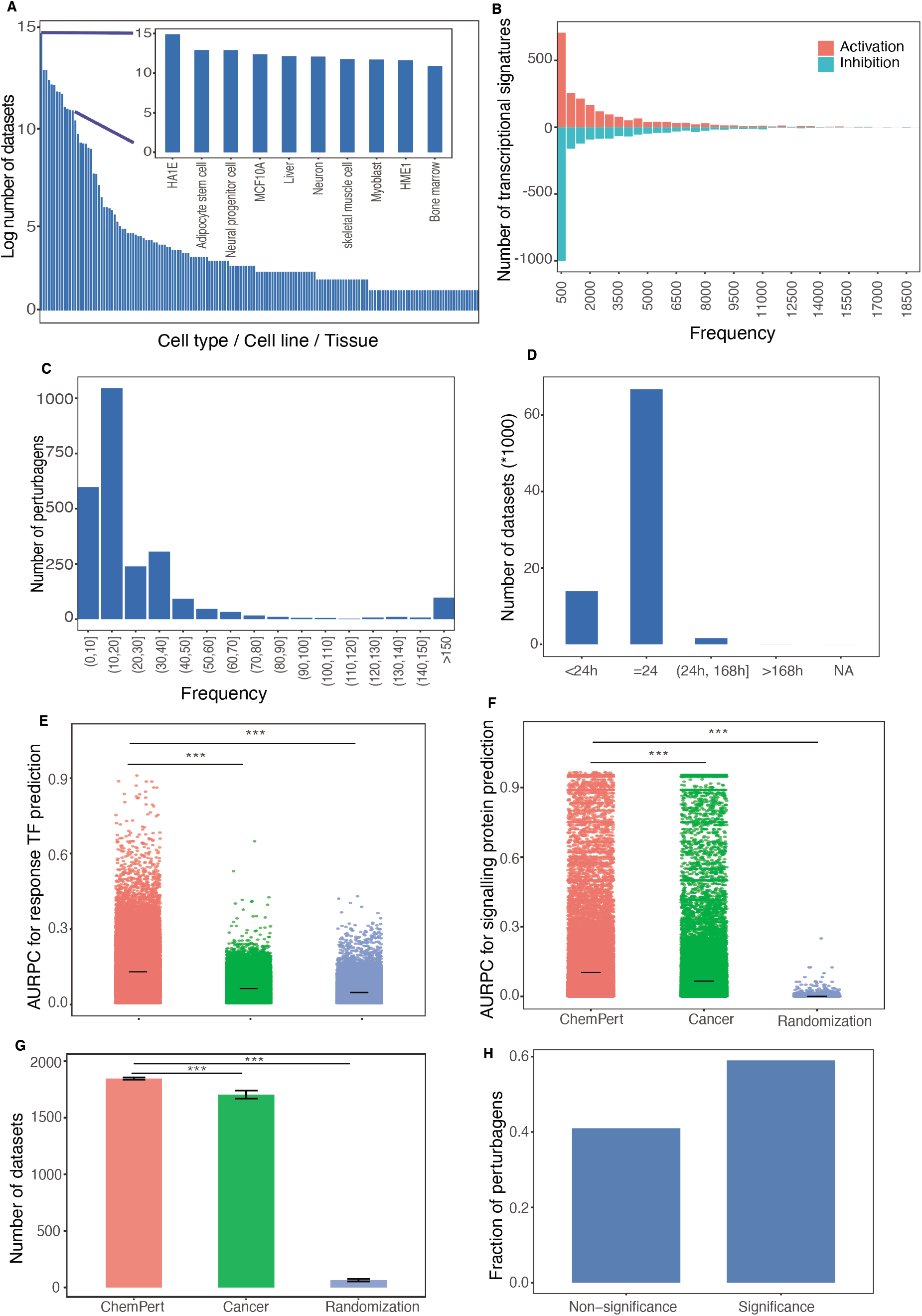
The compositions and evaluation of ChemPert database. (**A**) Distribution of datasets across different cell types/lines/tissues in the ChemPert database. Y-axis scale is log2(number+1) for each cell type/line/tissue. (**B**) Frequency of TFs in the ChemPert database, including inhibited and activated ones. X-axis represents the frequency of TFs and y-axis presents the number of TFs with corresponding frequency. (**C**) Distribution of perturbagen frequency in the ChemPert database. X-axis represents the frequency of perturbagen, and y-axis represents the number of perturbagen with corresponding frequency. (**D**) Distribution of datasets for different perturbation durations. (**E**) AURPC for response TF prediction given perturbagens. (**F**) AURPC for protein target prediction given response TFs. (**G**) Number of datasets with correct perturbagen prediction, data are *mean* ± *MSE*. E-G used the benchmarking datasets to compare the performance of ChemPert tool using the ChemPert database, cancer database or randomization. Significance was calculated by using one-sided Wilcoxon test. ***: *P-value <2.22e-16*. (**H**) Fraction of perturbagens whose within-ness are significantly larger than between-ness.

### Benchmarking of ChemPert

The mapping between signalling perturbations and response TFs enables *in silico* predictions of either the downstream effects of given perturbagens or the perturbagens that can target given sets of TFs. In particular, such mapping for non-cancer cells will significantly reduce our efforts for identifying perturbagens of desired effects instead of the perturbagens killing cells in cancer therapies, which will aid in a wide range of biological and clinical applications. Therefore, we developed a computational tool for either predicting downstream response TFs given a perturbagen of known target proteins, or the perturbagens of desired TF responses.

To evaluate the importance of using the ChemPert database, rather than cancer cell databases, for the prediction in non-cancer cell types, we conducted a benchmark analysis on the ChemPert database and on the cancer database solely consisting of cancer perturbation datasets (see Methods). The results show a significantly higher performance (measured as the area under precision-recall curve (AUPRC)) with the ChemPert database than with the cancer database in the prediction of response TFs (Figure 2E). In fact, the perfomane of the latter was similar to the random selection of reference datasets (Figure 2E). We also investigated if a similar predictive performance could be achieved without taking into account the initial cell states (i.e., based only on perturbagen target similarities). This result shows a significant decrease in the performance (Figure S2A), indicating that perturbagen similarity alone is not sufficient for mapping cell-specific response TFs.

As for the prediction of perturbagens from response TFs, the AUPRC of signalling protein targets was significantly better when using the ChemPert database compared to using the cancer database (Figure 2F). Moreover, using the ChemPert database significantly increased the number of datasets with true perturbagen prediction (Figure 2G) and the rank of true perturbagens was significantly lower (Figure S1B). To further investigate why the cancer database showed a worse performance, we identified the perturbagens commonly used in both ChemPert and cancer databases and found that around 60% of the perturbagens had significantly larger within-ness than between-ness (see Methods). This result indicates that the rewiring of upstream signaling pathways in cancer cells could induce significant changes in downstream TF responses compared to non-cancer cells.

Taken together, these results highlight the importance of use of non-cancer cell perturbation datasets, rather than cancer-based datasets such as Cmap and LINCS 1000, for mapping between signalling perturbations and response TFs in non-cancer cells. Indeed, ChemPert is able to make more accurate predictions by taking advantage of the non-cancer cell database.

### Description of ChemPert web interface

The ChemPert web interface mainly includes two sections (Figure 3A): the database (Figure 3B) and the webtool (Figure 3C). The database section allows users to browse, search and download any datasets in ChemPert without creating an account and login. The home page of the database section provides a summary of the database and allows users to get access to one of the three main resources of the databases, the targets of perturbagens, the gene expression profiles of initial cellular states and the TF responses after perturbations (Figure 3B). For example, when users click the button “Transcriptional responses”, a table listing the major meta information on each dataset will be returned, including the perturbagen, data accession number, cell type, perturbation duration and concentration (Figure 3D). The search area allows users to search for the datasets of interest based on the perturbagens, cell types or species (Figure 3D). In particular, users can click the “Response ID” to browse the response TFs of corresponding dataset (Figure 3E). Clicking the “Perturbagen” button enables the users to browse the protein targets of this chemical compound (Figure 3F). In addition, users can download the datasets of interest or download all datasets from “Download” page.

**Figure 3.**
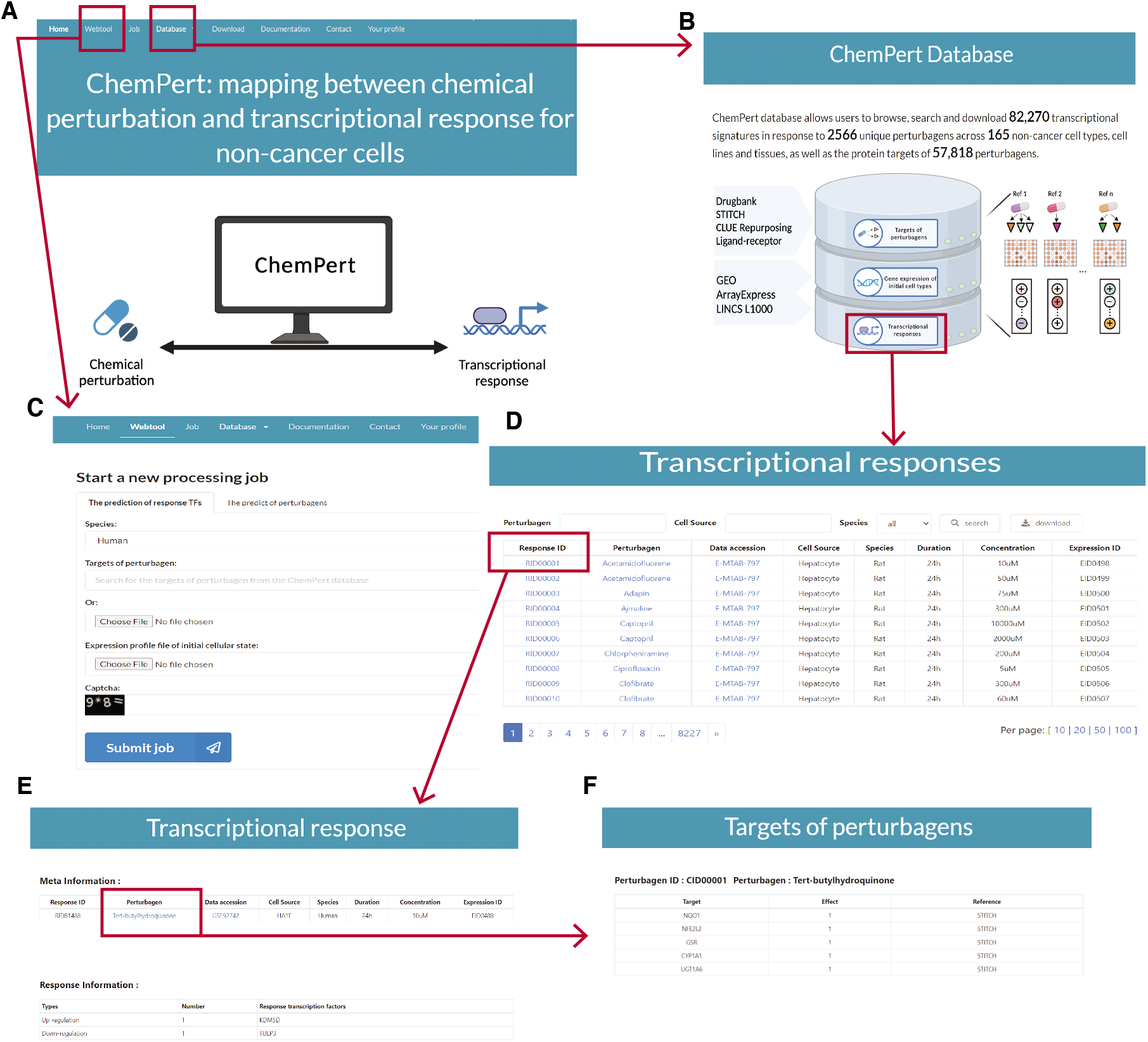
Illustration of ChemPert web interface. (**A**) The home page of web interface. ChemPert mainly consists of two sections: database and webtool. (**B**) The home page of the ChemPert database. The database is composed of three parts: targets of perturbagens, gene expression of initial cell types and transcriptional response. Clicking the button can switch to corresponding part. (**C**) The webtool page. Users can predict either the response TFs of given perturbagen or the perturbagens targeting desired query TFs. (**D**) The transcriptional response table listing the meta information of datasets. (**E**) Detailed transcriptional response for one dataset. (**F**) Information about targets of perturbagens.

The webtool section provides an intuitive interface for users to predict either response TFs or perturbagens (Figure 3C). To predict response TFs of a query perturbagen, users can search for the targets of perturbagen in the ChemPert database as input. If a query perturbagen is not available in the database for the prediction of response TFs, users can still run the tool by providing the protein targets of the query perturbagen as input. After providing the required input, users can track the status of the job and download the final predictions in “Jobs” page. The detailed usage of ChemPert web interface is described in “Documentation” page.

### Use case - ChemPert predicts cell state-specific responses to drugs in NASH

NASH is an advanced form of non-alcoholic fatty liver disease that can be lethal and currently no FDA-approved medications exist (6,7). We applied ChemPert to the RNA-seq data of two models of diet-induced NASH to predict the TF responses of perturbagens that could enable us to find optimal treatments. The first model consists of mice fed with a high-fat diet rich in fructose, palmitate, and cholesterol for 20 weeks (FCP) (8). The second model consists of mice fed with a choline-deficient, methionine-reduced high-fat diet for seven weeks (CDA) (9). In addition, both models were stratified into two groups based on the severity of the liver disease phenotype: mild NASH and advanced NASH. Mice with advanced NASH had significantly more inflammatory foci and collagen fiber formation compared to mice with mild NASH (10). The use of both diet models and their two disease severity phenotypes allows us to take advantage of the heterogeneous NASH states and make more reliable assessment of predicted response TFs, as an effective drug for the treatment of NAFLD must be effective at different stages. ChemPert was run for three perturbagens: obeticholic acid (OCA) known to significantly improve fibrosis in adult patients with definite NASH (11); pioglitazone and vitamin E, associated with reductions in hepatic steatosis and lobular inflammation, but with no improvement in fibrosis score (12).

In the case of OCA, 209 TFs were predicted to be upregulated in the CDA model, 135 of which were predicted to be overexpressed in both mild and severe models (Figure 4A). In the FCP model, upregulation of 203 TFs in response to OCA was predicted regardless of disease severity. Among all these TFs, 40 were common in both NASH models. Due to the low number of common TFs, the GSEA analysis did not identify any enriched pathway. However, consistent with the recognized therapeutic effect of OCA, these common TFs are related not only to hepatic steatosis and steatohepatitis improvements (ATF6, HBP1, BTG1, SAP18, PPARD, PPARG, BIRC2), but also to anti-fibrotic effects (FOXO1, INSR, KLF6) and blocking of disease progression (DACH1, RYBP, ZFP36L1). Similarly, the 42 common downregulated TFs (Figure 4B) include both signatures of steatosis and obesity (CNOT3, CREB3L3, REPIN1, STAT1), and signatures of fibrosis (CCNE1, ETS1, HDAC6, HDAC9, HLF, PLAGL1, SOX4, TRIM16, TRIM29) and hepatocellular carcinoma (HCC) (BCL3, MYCBP, SMARCA4). The detailed explanation for each TF can be found in Supplementary note.

**Figure 4.**
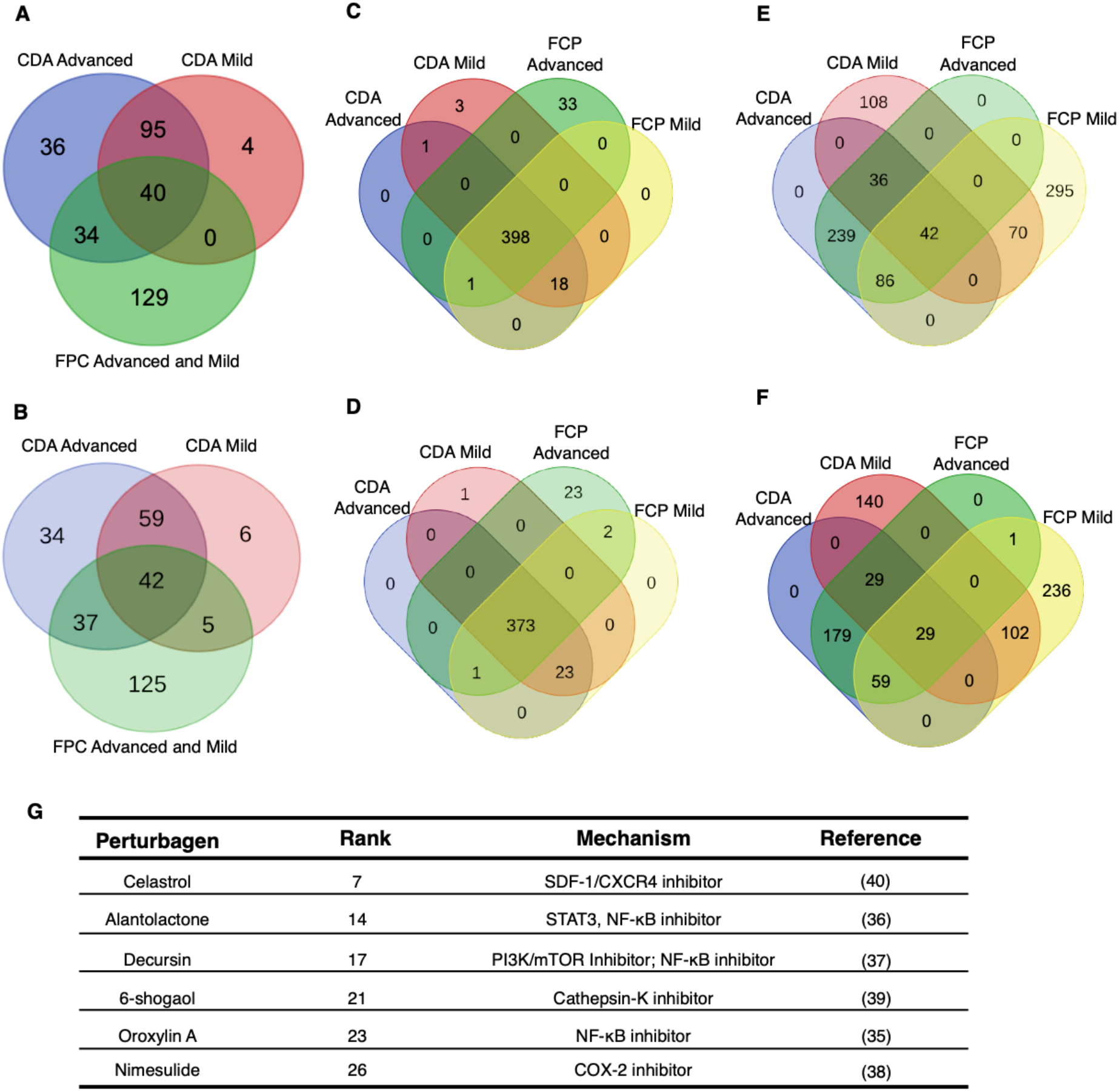
Application of ChemPert. Venn diagrams showing overlaps of predicted TFs among different diets and disease states of NASH models. Up-regulated TFs (**A**) and down-regulated TFs (**B**) after OCA perturbation, up-regulated TFs (**C**) and down-regulated TFs (**D**) after pioglitazone perturbation, up-regulated TFs (**E**) and down-regulated TFs (**F**) after vitamin E perturbation. (**G**) The representative of predicted perturbagens with literatures support for the treatment of OA. Details are shown in Table S4.

The pioglitazone perturbation predicted 421 and 449 total up-regulated TFs in the CDA and FCP models, respectively, 398 of which are common to both disease models and disease states (Figure 4C). The GSEA of these 398 TFs (Table S1) contained *Nuclear Receptor transcription pathway* including PPARD and PPARG, as expected, since the thiazolidinediones, such as pioglitazone, are synthetic agonists for these receptors, that play a key role in lipid metabolism. However, the GSEA also produced *TGF-β signaling* which is a well-known profibrogenic cytokine due to its role in hepatic stellate cell (HSC) activation and extracellular matrix production. This pathway has been described to contribute to all stages of liver disease progression, from initial liver injury through inflammation and fibrosis to cirrhosis and hepatocellular carcinoma (HCC) (13–15). Moreover, *TRAF6 Mediated Induction of proinflammatory cytokines* is a key driving force of proinflammatory and profibrogenic responses in NASH (16) and has been described as a possible contributor to progression to HCC (17). *TLR4 signaling repertoire* is involved in a variety of liver injury including that induced by NASH, which has been shown to play a key role during fibrogenesis in preclinical models of NAFLD (18), as wells as to enhance TGF-β signaling (19). *Stabilization of p53* has also been involved in the pathogenesis of fatty liver disease (20). On the other hand, the GSEA of 376 common down-regulated TFs (Figure 4D, Table S2) included the *Interferon gamma* (IFN-γ) *signaling*, which has previously shown promising results in terms of fibrosis scores in patients with chronic HBV infection, most likely by antagonizing profibrogenic transforming TFG-β effects (21); and in accordance with these data, a preclinical IFN-γ deficient model showed a rapid development of liver fibrosis when fed a fatty diet (22).

Finally, the vitamin E perturbation obtained 581 and 768 total upregulated TFs for the CDA and FCP models, respectively, 42 of which are common to both disease models and disease states (Figure 4E). The GSEA of these TFs (Table S3) identified, as in Pioglitazone, the *Nuclear Receptor transcription pathway*, but also the *Regulation of Lipid Metabolism by Peroxisome proliferator-activated receptor alpha* (*PPAR-α*). Furthermore, the *Toll Like Receptor 3 (TLR3) Cascade* and *TRIF mediated TLR3 signaling* were enriched. Activation of TLR3 in HSCs has been demonstrated to exacerbate liver fibrosis (23). The GSEA of 29 common down-regulated TFs (Figure 4F) did not result in any enrichment. However, these TFs include FOXO1 and KLF6, which identified as anti-fibrotic (24–26) that were predicted to be upregulated in the OCA perturbation. Others are ID2, which reduces differentiation of HSCs and thus inhibits liver fibrosis (27), RUNX1, which regulates the expression of angiogenic and adhesion molecules, enhancing inflammation and disease severity in NASH (28), and KLF2, which has been reported to be elevated in livers from obese mice, and to induce triglycerides accumulation (29).

Overall, the analysis with OCA predicted the up-regulation of TFs related to the inhibition of HSC activation responsible for the collagen deposition in liver tissue during fibrogenesis (30), along with TFs described as protective against inflammatory response and hepatic fat deposition, and down-regulation of TF signatures of steatosis, fibrosis and HCC. Although the common TFs of pioglitazone and vitamin E perturbations appeared to be viable for treating hepatic steatosis and inflammation, none of these were associated with improvement of fibrosis. Thus, this analysis demonstrates that ChemPert is valid for predicting the transcriptional effects of different drugs at different stages of NAFLD and could be a useful tool for pre-screening a wide range of chemical treatments prior to the pre-clinical or clinical studies.

### Use case - ChemPert predicts novel perturbagens for the treatment of OA

OA is a complex degenerative disease leading to disability and characterized by cartilage degradation, synovial inflammation, and bone remodelling (31). Currently, effective pharmacologic therapies for OA are still not available and more specific approaches are desirable (32). Thus, ChemPert was applied to OA to investigate potential therapeutic treatments. The differentially expressed TFs in human osteoarthritis cartilage compared to non-osteoarthritis individuals were identified as input (GSE169077). A considerable number of known clinical or pre-clinical chemical compounds for the treatment of OA were recapitulated by ChemPert and ranked among the top 30 (Figure 4G, Table S4). The nuclear factor-kappaB (NF-κB) signaling pathway is regarded as potential targets for the therapeutic treatment of OA, since NF-kB is aberrantly upregulated in OA patients and NF-κB is included in many OA-associated events, including chondrocyte catabolism, chondrocyte survival, and synovial inflammation (33,34). In agreement with this, several perturbagens targeting NF-κB were predicted by ChemPert, including oroxylin A (35), alantolactone (36) and decursin (37), which all have been shown to ameliorate OA. These perturbagens attenuate OA progression by inhibition of inflammatory response, hypertrophy, cartilage degeneration or impaired autophagy triggered by IL-1β. Moreover, ChemPert also predicted the perturbagen, nimesulide, a cyclo-oxygenase (COX)-2-selective inhibitor that attenuates the pain associated with walking for OA patients (38). The prediction 6-shogaol has been shown to significantly reduce the hypertrophic markers in cartilage and prevent synovial inflammation and cartilage degradation in OA (39). Celastrol was also predicted, which is known to target SDF-1/CXCR4 signalling pathway is able to attenuate pain and cartilage damage in OA (40) and has the potential to prevent OA by inhibiting the ERs-mediated apoptosis (38). Studies also revealed that the PI3K/AKT/mTOR pathway plays a crucial role in cartilage degradation and can be used as a therapeutic target for the clinical intervention of OA (41,42). Consistently, we identified the signaling proteins that are enriched in PI3K/AKT pathway (Figure S2) and the perturbagens that inhibit the PI3K/AKT signaling pathway, including oroxylin A (43), KU-0063794 (44), and other novel perturbagens such as NVP-BEZ235 and TG100-115 (Table S4). In addition, previous reports have indicated that VEGF can be a biomarker for patients with OA, which is highly expressed in articular cartilage, synovium, subchondral bone and serum of OA patients (45). Indeed, we identified the signaling proteins that are enriched in VEGF pathway and predicted corresponding inhibitors, like WHI-P180 and PP-121. Furthermore, another novel prediction is 1,5-isoquinolinediol, a PARP-1 inhibitor. In accordance with our prediction, a previous study also reported that PARP-1 inhibitors are able to decrease the inflammatory response in the cartilage of OA rat model (46).

To summarize, ChemPert not only recapitulated the known perturbagens, but also provided novel predictions as potential therapies for the treatment of OA. These results demonstrate the usability of ChemPert for *in silico* chemical screening and drug discovery, and can be generally applicable to different diseases to prioritize the perturbagens that reverse the disease phenotypes to the healthy counterparts.

## Methods

### Construction of ChemPert database

In this study, we constructed a database depicting the relationship between chemical perturbations, protein targets of perturbations and downstream transcriptional signatures. We considered the responses of transcriptional regulators including transcription factors, transcriptional co-factors and chromatin remodelling factor as transcriptional signatures and used the term ‘response TFs’ to refer to these signatures for brevity. First, we collected transcriptome profiles of chemical perturbations (including small molecules, growth factors, cytokines and other protein ligands) from Gene Expression Omnibus (GEO) (47) and ArrayExpress (48). Specifically, the keywords commonly used in perturbation studies, such as ‘time series’, ‘response’, ‘treat’, ‘perturb’, ‘presence’ and ‘effect’, were used to search for the datasets in GEO and ArrayExpress. Then, we manually curated the datasets focusing on noncancer cell types/lines or tissues in human, mouse and rat (Figure 1A). The datasets were preprocessed, including background correction and normalization, either with the same approaches from the original studies or using the limma R package (v3.38.3) (49). In addition, we also extracted the chemical perturbation datasets of non-cancer cells from LINCS L1000 at Level 3, where the quantile normalization was performed (2). The response TFs of each perturbagen were obtained by performing differential expression analysis using the limma R package. The genes with Benjamini-Hochberg (BH) adjusted p-value <= 0.05 and absolute fold change >=1.5 were considered as differentially expressed genes (DEGs) compared to unperturbed control samples when the sample replicates were larger than two. Otherwise, only the fold change was used as the criterion. Differentially expressed TFs were considered as response TFs and were identified from DEGs based on AnimalTFDB 2.0 (http://bioinfo.life.hust.edu.cn/AnimalTFDB2/) (50), which contains the information of transcription factors, transcriptional co-factors and chromatin remodelling factors. Furthermore, these response TFs were assigned with Boolean value 1 and −1, which represented up-regulation and down-regulation after perturbation, respectively. The gene symbols of mouse and rat were converted to human orthologue gene symbols with the Biomart R package (v2.38.0) (51). The gene expression profile of each dataset before perturbation was denoted as an initial gene expression profile (Figure 1A).

In addition, the direct signalling protein targets of perturbagens were retrieved from Drug Repurposing Hub (www.broadinstitute.org/repurposing) (52), DrugBank (www.drugbank.ca) (53), and STITCH v5.0 (http://stitch.embl.de) (54) (Figure 1A). In STITCH, only the targets with a confidence value larger than 0.4 were kept along with the experiment and database evidence. The receptor targets of protein ligands were identified from manually curated ligand-receptor pairs from Ramilowski et al. (55). The effects of perturbagens on protein targets, activation, inhibition and unknown, were assigned with value 1, −1 and 2 respectively.

### Prediction of perturbation response TFs

The ChemPert tool for the prediction of response TFs after a query perturbation consists of three major steps (Figure 1B). In short, it first identifies TF response datasets perturbed with similar perturbagens as the query perturbagen. Then, it filters out the TF response datasets whose initial cell states are not similar to the query initial cell state. Finally, low frequency TFs in the retained TF response datasets are discard. The details are described below.

Step 1: A modified Jaccard similarity between a query perturbagen and reference perturbagens in the ChemPert database is computed by

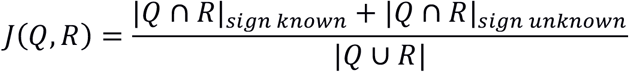

where |*Q* ∩ *R*|*_sign known_* is the cardinality of common protein targets with the same effect (activation or inhibition) between the query and reference perturbagens, whereas |*Q* ∩ *R*|*_sign unknown_* is the same cardinality computed among protein targets whose effects are unknown for the query and/or reference perturbagens. For the latter cardinality, a query protein target and a reference protein target are considered as a match as long as they both target the same protein regardless of the effect. Reference perturbagens with the modified Jaccard similarity higher than 1.5 z-score are retained. Then, all reference datasets perturbed by the retained perturbagens are retrieved from the ChemPert database.
Step 2: We reasoned that if the state of molecular paths from proteins targeted by a perturbagen to downstream TFs is similar between the query and reference datasets, the TF response of the query data will also be similar to the reference response TFs. To compute such similarity, the prior knowledge network (PKN) is constructed by merging ReactomeFI (56), Omnipath (57) and DoRothEA v2 (58). Then, the short paths from one signalling protein to each downstream TF are identified as follows: First, the shortest path lengths from each signalling protein to all downstream TFs are calculated using the unweighted breadth-first algorithm implemented in R package igraph. Subsequently, the path length that can reach the largest number of downstream TFs from that signalling protein is considered as the maximum path length. We then calculate all possible short paths between the signalling proteins and all downstream TFs that are within this maximum path length. This procedure is repeated for every signalling protein in the PKN. Then, for each signalling protein-TF pair, a path enrichment analysis is performed using Fisher’s exact test:

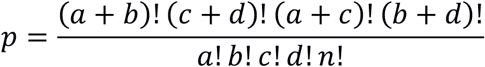 Where *a* is the sum of normalized gene expression values of proteins present in all the short paths including the starting signalling protein and target downstream TF, *b* is the sum of normalized gene expression values of all genes in the dataset, *c* is the number of proteins present in all the short paths, *d* is the total number of genes in the dataset, and *n* is the sum of *a, b, c* and *d*. The gene expression is normalized by the highest expression value in the dataset. Since Fisher’s exact test can accept only integer values, the decimal values are rounded for *d* and *c*. The p-values are corrected by the Benjamini-Hochberg method and paths with the adjusted p-value <= 0.05 are considered enriched. The initial cell state similarity between a query and a reference dataset is computed by the Jaccard similarity of common enriched paths. Reference datasets with this Jaccard similarity higher than z-score 1.5 are retained for the next step.
Step 3: The frequency of each response TF is computed among the reference datasets retained after Step 2. When a TF has both directions (i.e., up- or down-regulated), the one with the lower frequency is discarded. If this frequency is the same, the TF is discarded due to the uncertainty of its direction. Thus, the final output contains predicted response TFs in one direction and their frequency in the retained reference datasets.

### Prediction of perturbagens targeting query TFs

Given a set of query TFs, ChemPert is also available for the prediction of perturbagens. The tool first identifies the potential signalling protein targets from the ChemPert database whose perturbation can induce a similar set of response TFs. Then, the perturbagens whose protein targets are enriched among the predicted signalling proteins are further identified (Figure 1B). The similarity between query TFs and response TFs of each reference dataset in the ChemPert database is calculated by using a modified Jaccard similarity as:

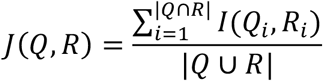

with indicator function:

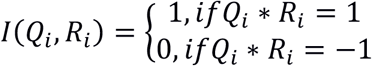

where *Q* is the set of query TFs and *R* is the response TFs for each reference in the ChemPert database. In order to ensure the consistent effect of a TF between the query and the reference, we modified the Jaccard similarity by adding an indicator function. If the TF has the same effect (both inhibition/activation), then 1 is assigned, and 0 otherwise. The perturbagens of the reference datasets are ranked based on the similarity in descending order. Only the highly confident perturbagens with z-score of similarity larger than 3.5 are selected for the further analysis. Next, ChemPert retrieves the signalling protein targets of each selected perturbagen from the ChemPert database and order the signalling proteins based on the sum of the similarity score of their corresponding perturbagens. The effects of signalling proteins are reported based on the majority effects of their perturbagens. For example, value 1 is assigned to the signaling protein when more predicted perturbagens have activation effects on it. The signaling protein is assigned as 2 when all of its predicted perturbagens have unknown effects on it.

Finally, the prediction of perturbagens is conducted as follows: Each perturbagen and corresponding protein targets in ChemPert database is converted into a regulon-like class as TF-regulons in database DoRothEA v2. The analytic Rank-based Enrichment Analysis (aREA) was carried out to identify the perturbagens whose protein targets are enriched in the top ranked predicted signaling proteins by using msviper function of Viper package (v1.18.1) (59). The predicted perturbagens are ranked based on the normalized enriched score (NES) and the ones with false discovery rate less than 0.05 are kept. The top 30 predictions are recommended as final predictions.

### Evaluation of ChemPert database

The predictive performance of the ChemPert database was compared to a cancer database using the subset of the LINCS L1000 database, which only contains cancer cell datasets (2). We performed a leave-one-out validation, in which one reference dataset in the ChemPert database was randomly selected as a query dataset and removed from database. This query dataset was used to compare the performance between using the ChemPert database and using the cancer database in terms of response TFs prediction and perturbagens prediction. This validation was performed by randomly selecting 4000 datasets and this procedure was repeated 10 times. In addition, the difference in transcriptional responses between noncancer cells and cancer cells was quantified using perturbagens that are commonly used for at least three cell types in both ChemPert and cancer database. The Jaccard similarity of transcriptional responses within non-cancer cells (within-ness) and that between non-cancer and cancer cells (between-ness) were calculated and compared. The perturbagens whose within-ness are significantly larger than the between-ness were identified by using one-side Wilcoxon test with adjust p-value < 0.05.

### Construction of ChemPert web interface

The ChemPert web interface was implemented using Python 3.7 (https://www.python.org/) programming language and constructed using the Django (https://www.djangoproject.com/), a high-level Python web framework. In the Django web framework, the front-end responsive web pages were built using the HTML templates combined with Semantic UI (https://semantic-ui.com/) and Bootstrap (https://getbootstrap.com/) libraries. The responsive table widget with filter, search and pagination functionalities in some web pages was implemented using django-filter (https://django-filter.readthedocs.io/) and django-tables2 (http://django-tables2.readthedocs.io/) libraries. The Django framework provides data-model syntax, the data is defined in the Django model and is easily mapped to the SQLite Database (https://www.sqlite.org/index.html). Finally, this web project was hosted on a Rocky Linux 8 (https://rockylinux.org/) server.

## Discussion

ChemPert provides the first comprehensive compendium of manually curated perturbation transcriptomics exclusively for non-cancer cells. In addition, ChemPert provides a computational tool that leverages the non-cancer cell data to predict either TF responses after perturbations, or perturbagens that target desired sets of TFs. Importantly, predictions generated for non-cancer cells when using ChemPert database were significantly more accurate than those based on cancer databases. ChemPert is freely available as a web application and will facilitate future *in silico* chemical compound screening for non-cancer cells and drug discovery for non-cancer diseases.

Due to the scarcity of available combinatorial perturbation datasets, we focus on transcriptional signatures of single-agent perturbations in the current version of ChemPert. However, our future plan is to continue adding new non-cancer combinatorial perturbation datasets to address the important challenge of in silico combinatorial drug screening. In addition, we will regularly collect and compile new single-agent perturbation datasets to maintain the state-of-the-art of the database.

## Data access

The ChemPert web interface is freely accessible at: https://chempert.uni.lu/. ChemPert was implemented in R and is available from Gitlab (https://git-r3lab.uni.lu/menglin.zheng/chempert).

## Acknowledgments

We thank Jacek Lebioda for the help of the development of web application.

## Contributions

A.d.S. conceived and supervised the overall study. M.Z. constructed and designed the database, developed the perturbagen prediction part of the tool and performed the analysis S.O. helped with the database construction, developed the TF response prediction part of the tool and performed the analysis. M.B. and M.M. gave an expert interpretation of the analysis of the NASH data. F.C. developed the database interface. M.Z., S.O., M.B., M.M. and A.d.S. wrote the manuscript.

